# Female rats self-administer heroin by vapor inhalation

**DOI:** 10.1101/2020.03.30.016725

**Authors:** Arnold Gutierrez, Jacques D. Nguyen, Kevin M. Creehan, Michael A. Taffe

## Abstract

Over the last two decades the United States has experienced a significant increase in the medical and non-medical use of opioid drugs, resulting in record numbers of opioid-related overdoses and deaths. There was an initial increase in non-medical use of prescription opioids around 2002, followed later by increased heroin use and then most recently fentanyl. Inhalation is a common route of administration for opioids, with a documented history spanning back to Mediterranean antiquity and up through modern use with e-cigarette devices. Unfortunately, preclinical studies using inhalation as the route of administration remain relatively few. This study was conducted to determine the efficacy of e-cigarette vapor inhalation of heroin in rats. Non-contingent exposure to heroin or methadone vapor produced anti-nociceptive efficacy in male and female rats. Female rats were trained to self-administer heroin vapor; the most-preferring half of the distribution obtained more vapor reinforcers when the concentration of heroin was reduced in the vapor vehicle and when pre-treated with the opioid receptor antagonist naloxone. The anti-nociceptive effect of heroin self-administered by vapor was identical in magnitude to that produced by intravenous self-administration. Finally, anxiety-like behavior increased 24-48 hours after last heroin vapor access, consistent with withdrawal signs observed after intravenous self-administration. In sum, these studies show that rewarding and anti-nociceptive effects of heroin are produced in rats by vapor inhalation using e-cigarette technology. Importantly, self-administration models by this route can be deployed to determine health effects of inhaled heroin or other opioids.

## 1. Introduction

Over the past two decades the United States has experienced a significant increase in the problematic use of opioid drugs. The trend started with prescription opioids circa 2002, followed by an increase in the use of heroin starting around 2012, and eventually added illicit fentanyl (and derivatives) starting in 2015; this all resulted in record numbers of opioid-related overdoses and deaths (Hedegaard et al., 2018; Rudd et al., 2016a; Rudd et al., 2016b; Scholl et al., 2018). Individual progression from the initial use of prescription drugs into illicit opioids often results in a change from oral ingestion to intravenous injection or inhalation of the volatilized drug (Mars et al., 2014; Woodcock et al., 2015). While the effects of injected opioids in humans and animals have been studied extensively, study of the effects of inhaled drugs in animals have been much less common, partly due to the lack of effective means of drug delivery for common laboratory species such as rats or mice. The recent adaptation of popular Electronic Drug Delivery Systems (EDDS; aka “e-cigarettes”) for research purposes has greatly facilitated the study of effects that drugs produce via inhalation.

Inhalation has been a common route of administration for opioid drugs, with a documented history spanning back to Mediterranean antiquity, as reviewed (Kritikos, 1960). Some of the earlier modern Western opioid misuse crises occurred in the 19^th^ century during an interval of Chinese diaspora and primarily involved the inhalation of opium vapor (Kane, 1881). Inhalation continues to be a common route of administration among individuals that abuse opioids at present (Alambyan et al., 2018) and some newer users will first choose inhalation as a means of administering opioids because of the perceived safety compared with injection (Stover and Schaffer, 2014). Although some degree of increased safety may be achieved, such as in the prevention of bloodborne pathogen transmission, inhalation and injection similarly facilitate progression into dependence (Barrio et al., 2001). Considering that the subjective and physiological effects of inhaled heroin are similar to those of injected heroin, but plasma levels may decline more quickly (Jenkins et al., 1994); this route might support more frequent re-dosing. Inhalation of opioids is often achieved by heating the drug over a surface and inhaling the resulting vapors (Jenkins et al., 1994; Strang et al., 1997). Recently, however, there have been reports of EDDS usage (vaping) for administering illicit substances including cannabis extracts and, most pertinently, opioid drugs (Blundell et al., 2018; Breitbarth et al., 2018; Kenne et al., 2017). The current popularity and acceptability of vaping, the covert and easily concealed designs of newer vaping devices, and the myriad flavorant options, which can potentially mask public use, make this latter method especially concerning. Because non-medical use of opioids by inhalation remains prevalent, further determination of any route-specific effects warrants attention.

Despite numerous epidemiological and clinical studies on the use of inhaled opioid drugs, preclinical reports which use inhalation as the route of administration remain relatively scarce. A limited literature shows that antinociceptive effects, as measured by the tail-withdrawal test, can be produced following passive exposure to vaporized morphine, heroin, and fentanyl in mice (Lichtman et al., 1996) and oxycodone in rats (Nguyen et al., 2019). Additionally, rhesus monkeys will self-administer heroin by inhalation and will exhibit behaviors that are characteristic of heroin intravenous self-administration, such as drug-seeking upon increase of the reinforcement contingency and dose-dependent suppression of drug-taking behavior when pre-treated with naloxone (Mattox and Carroll, 1996; Mattox et al., 1997). Adult rats will self-administer the synthetic opioid sufentanil in aerosol mist form delivered through a nebulizer (Jaffe et al., 1989) and rats and mice will self-administer sufentanil and fentanyl, respectively, through the use of a vapor delivery system based on EDDS technology (Moussawi et al., 2020; Vendruscolo et al., 2018). The opioid self-administration studies found behavioral effects comparable to those seen in rodent intravenous self-administration studies, such as concentration-dependent responding for drug, and the Vendruscolo et al. (2018) study additionally demonstrated escalation after extended drug access and enhanced drug-seeking behavior in naloxone pre-treated rats (Jaffe et al., 1989; Vendruscolo et al., 2018). Together, these studies demonstrate the viability of self-administration models of inhaled drugs and illustrate some of the behavioral similarities between the effects produced by inhalation and intravenous injection of opioid drugs. However, the limited number of available reports on the behavioral effects of opioid inhalation makes any conclusions on behavioral similarities or differences due to route of administration difficult to make. Further work is therefore required to determine the specific reinforcing properties of individual opioid drugs delivered by EDDS and other devices, especially under differing techniques for drug volatilization (see (Miliano et al., 2020; Moore et al., 2020) for review).

The present study was therefore designed to assess the effects of inhaled heroin vapor delivered by EDDS on measures of reinforcement, withdrawal-like behavior, and nociception. The approach adapted a vapor inhalation technique we have shown delivers behaviorally active, non-contingent doses of drugs as varied as methamphetamine and MDPV (Nguyen et al., 2016a; Nguyen et al., 2017), Δ^9^-tetrahydrocannabinol and cannabidiol (Javadi-Paydar et al., 2019a; Javadi-Paydar et al., 2018; Nguyen et al., 2016b), oxycodone, heroin and sufentanil, (Gutierrez et al., 2020a; Nguyen et al., 2019; Vendruscolo et al., 2018) and nicotine (Javadi-Paydar et al., 2019b). To determine behavioral efficacy of inhaled heroin, we first assessed anti-nociception by measuring tail-withdrawal latency from a hot water bath after non-contingent opioid vapor exposure and compared this with the effects of subcutaneous opioid injection. Next, groups of adult female Wistar rats were trained to lever-press for delivery of in-chamber heroin vapor. Female rats were selected for this study first because the prior inhalation studies used male rats (Jaffe et al., 1989; Vendruscolo et al., 2018) and it is important to determine if effects generalize to female subjects (Cicero et al., 2003; Clayton and Collins, 2014; Shansky, 2018; Shansky and Woolley, 2016). Furthermore, there is some indication female rats more readily self-administer more morphine, heroin, oxycodone, or fentanyl compared with male rats (Cicero et al., 2003; Klein et al., 1997; Nguyen et al., 2020), thus female rats were anticipated to speed model development. Following acquisition of vapor self-administration, drug concentrations in vapor solution were varied and behavioral response was assessed. Drug-seeking behavior was also examined after the administration of the antagonist naloxone to provide converging evidence on the pharmacological specificity of the behavior. Further, the self-administration rats were evaluated for nociceptive responses before and after a self-administration session to further verify that an active dose was self-administered and to compare the magnitude with effects observed in a traditional intravenous self-administration paradigm. Finally, anxiety-like behavior was probed immediately, 24-, and 48-hours post-session using an elevated plus maze approach.

## 2. Material and methods

### 2.1 Animals

Adult female (N=8) and male (N=8) Wistar rats (Charles Rivers Laboratories) were used for non-contingent vapor and injection studies. Groups of adult female (N=16) Wistar rats (Charles Rivers Laboratories) were used for vapor self-administration studies. Rats used for non-contingent vapor and injection studies, and for vapor self-administration, began testing at 10-11 weeks of age. Additional groups of adult male (N=17) Wistar rats were used for a study of the anti-nociceptive effects of intravenously self-administered heroin. Rats used for this comparison study were 39-weeks old at the time of testing. Rats were acclimated to the facility’s vivarium upon arrival for two weeks prior to initiating studies. The vivarium was kept on a 12:12 hour light-dark cycle and all studies were conducted during the rats’ dark period. Food and water were provided *ad libitum* in the home cage and body weights were recorded weekly. Two experimental cohorts of female rats were used for vapor self-administration experiments (N=16 total). The first subgroup (N=8), designated Cohort 1 (C1), underwent acquisition training, nociception testing, and concentration substitution experiments (further described below). Animals in Cohort 2 (C2) were tested as described for C1 but were returned to self-administration prior to being challenged with naloxone to assess alterations in self-administration and then prior to being evaluated in an elevated plus maze (EPM) to assess anxiety-like behavior after spontaneous withdrawal as outlined below. Experimental procedures were conducted in accordance with protocols approved by the IACUC of The Scripps Research Institute and consistent with recommendations in the NIH Guide (Garber et al., 2011).

### 2.2 Drugs

Heroin (diamorphine HCl) was administered by s.c. injection and by vapor inhalation (see below). Methadone was administered by s.c. injection and naloxone by i.p. injection. Inhalation/vapor doses are varied, and are therefore described, by altering the concentration in the propylene glycol (PG) vehicle, the puff schedule and/or the duration of inhalation sessions in this approach (Javadi-Paydar et al., 2019b; Nguyen et al., 2016a; Nguyen et al., 2016b). Heroin was dissolved in PG to achieve target concentrations and then loaded in e-cigarette tanks in a volume of ∼ 0.5 ml per session. Fresh solutions were used for each session. Naloxone, methadone and heroin were dissolved in saline (0.9% NaCl), for injection. The heroin was provided by the U.S. National Institute on Drug Abuse and the PG, methadone and naloxone were obtained from Sigma-Aldrich Corporation (St. Louis, MO, USA).

### 2.3 Electronic Vapor Drug Delivery

Passive opioid vapor (heroin, methadone, or oxycodone; all 100 mg/mL in PG vehicle), exposure sessions prior to the initial nociception experiments were conducted in modified Allentown rat cages (259 mm X 234 mm X 209 mm) with the use of Model SSV-1 e-vape controllers (La Jolla Alcohol Research, Inc, La Jolla, CA) set to 5 watts to trigger Protank 3 Atomizers (Kanger Tech, Shenzhen Kanger Technology Co. LTD; Fuyong Town, Shenzhen, China) pre-filled with drug solution. A computerized controller (Control Cube 1; La Jolla Alcohol Research, Inc, La Jolla, CA, USA) was used to automatically trigger ten-second vapor puffs at five-minute intervals. Vacuum exhausts attached to the chambers pulled ambient air through intake valves at ∼2 L per minute, starting 30 seconds before each scheduled vapor delivery. Vacuum exhausts were closed immediately following each vapor delivery. Duration for passive vapor exposure sessions was 30 minutes.

For self-administration experiments, vapor (heroin 50 mg/mL in PG vehicle for acquisition and maintenance) was delivered into sealed vapor exposure chambers (152 mm W X 178 mm H X 330 mm L; La Jolla Alcohol Research, Inc, La Jolla, CA, USA) through the use of e-vape controllers (Model SSV-3; 58 watts; ∼214 °F, La Jolla Alcohol Research, Inc, La Jolla, CA, USA) to trigger Smok Baby Beast Brother TFV8 sub-ohm tanks. Tanks were equipped with V8 X-Baby M2 0.25 ohm coils. Vapor exposure chambers were attached to vacuum exhausts which continuously pulled ambient air through intake valves at ∼1.5-2 L per minute. Vapor deliveries were 1 second in duration for self-administration sessions, and drug vapor from each delivery was mixed with flowing air entering the chamber. These parameters resulted in a vapor dwell time of approximately 30 seconds, in those cases where no further vapor deliveries had been obtained. Sessions were generally run on sequential weekdays, except when holidays occurred.

### 2.4 Behavioral Responses to Vapor Inhalation of Heroin

#### 2.4.1 Nociception assays

Tail withdrawal assays were performed using a Branson Brainsonic CPXH Ultrasonic Bath (Danbury, CT) filled with water and set and maintained at a temperature of 52°C. The bath was stirred, and temperature was verified using a second thermometer prior to each measurement. A stopwatch was used to measure the latency to tail withdrawal, and a 15-second maximum time limit to withdraw was imposed for each session.

Three sets of studies were conducted to examine the anti-nociceptive effects of inhaled heroin vapor by measuring tail-withdrawal latency from a hot water bath. The first set of studies were conducted to verify vapor exposure parameters that would induce significant anti-nociception, to compare this to an injected dose commonly reported in precedent literature and to determine if effects generalized to another opioid, methadone. For this, groups (N=8) of male and female rats underwent 30-minute exposure sessions to PG vehicle, methadone (100 mg/mL), oxycodone (100 mg/mL), or heroin (100 mg/mL). Tail-withdrawal measurements were then performed immediately following a five-minute vapor clearance interval, and again at 60, 90, and 120 minutes after start of vapor exposure. Rats then received subcutaneous injections of saline, oxycodone (2, 4 mg/kg), methadone (4 mg/kg), or heroin (1 mg/kg). Tail-withdrawal measurements were again performed 35 minutes after injections and again at 60, 90, and 120 minutes post-injection. There was a three-day minimum interval between test sessions in these experiments. Experimenters performing the tail-withdrawal were blinded to treatment conditions.

Next, studies were conducted to determine the anti-nociceptive effects of inhaled heroin (50 mg/mL in PG vehicle) immediately following a one-hour (N=16) and a 30-minute (N=8) heroin vapor self-administration session in female rats. For self-administration rats, a pre-session baseline tail-withdrawal assessment was performed immediately before the first self-administration session, i.e., prior to any drug exposure. After commencement of self-administration training, a nociception assessment was performed prior to, and immediately after, the eighth one-hour session in both Cohorts. A second set of nociception assessments were performed on C2 prior to, and after, the 20^th^ session. The duration for this latter self-administration session was reduced to 30 minutes to constrain the variability in drug intake pattern and the corresponding variability in drug distribution and metabolism. Start times for self-administration sessions for each rat were offset by five minutes to allow for the time required to complete the tail-withdraw measurement for each rat. Experimenters performing the assay were blind to how many vapor deliveries a given rat had obtained.

Finally, the similarity in degree of anti-nociception between i.v. heroin and inhaled heroin vapor was assessed. To this end, tail-withdrawal assessments were performed before and after a 30-minute i.v. heroin (0.006 mg/kg/infusion) self-administration session in a separate group of male rats (n=16). This group consisted of male Wistar rats that had been exposed to PG vehicle vapor or 100 mg/mL THC in PG vapor during adolescence and trained as adults to intravenously self-administer oxycodone in 8 h session (Nguyen et al., 2020). They had been switched to heroin IVSA for this study. For this experiment, a pre-session tail-withdrawal assessment was performed which was followed by a 30-minute heroin i.v. self-administration session at a dose of 0.006 mg/kg/infusion. Withdrawal latencies were again measured immediately after the session. There were no main effects of the prior adolescent repeated THC exposure or interactions of adolescent exposure with the time of assessment confirmed, thus the data presented were collapsed across adolescent treatment groups.

#### 2.4.2 Self-Administration Acquisition

Training began one day after the two-week acclimation period. Behavioral testing was performed in a behavioral procedure room under red lighting. Vapor self-administration chambers were housed within an opaque black holding rack which remained closed throughout testing. Chambers were thoroughly cleaned, and bedding was replaced after each session. Upon initiation of a session, rats had access to two levers located on the right and left wall at the end of the chamber away from the door. Pressing the active drug-associated lever produced a one-second vapor delivery and illuminated a light which was located above the lever. Each delivery was followed by a 20-second time-out period during which the light remained on and the drug-paired lever became inactivated. Following the 20-second time-out period, the chamber light turned off and the drug-paired lever once again became active. Pressing the non-paired inactive lever at any time throughout the session was recorded but was without consequence. Any active-lever presses, including those occurring during time-out periods, were recorded. Animals initially underwent 10 one-hour FR (1) daily sessions with the drug at a concentration of 50 mg/mL heroin in the PG. The reinforcement contingency was then increased to an FR (2) schedule, which remained throughout the study, and all rats underwent three more one-hour sessions to complete acquisition.

Rats in cohort 1 (C1) completed the 13 self-administration training sessions and then testing in the concentration substitution experiments. Following acquisition, rats in C2 continued for six more one-hour sessions, one 30-minute session immediately before a nociception assessment, and seven two-hour sessions. Concentrations were then substituted over 45 two-hour sessions. During this time, heroin was delivered at a concentration of 25 mg/mL across eight sessions and at 1 mg/mL over six. PG vehicle was given over the next 13 sessions, and the concentration was then increased back to 50 mg/mL for six sessions. Separate THC-heroin experiments were conducted across the following 12 sessions (data not shown). There was a four-day washout period before rats returned to the 50 mg/mL heroin training concentration for the EPM testing phase. Animals then proceeded to concentration substitution experiments followed by challenge with the antagonist naloxone.

#### 2.4.3 Concentration substitution

Following acquisition training, rats in C1 underwent two-hour sessions on an FR (2) reinforcement contingency during which drug concentrations were altered (1 mg/mL, 50 mg/mL, and 100 mg/mL) for each session, in a counter-balanced order with every rat exposed to each concentration once. Animals in C2 were assessed for concentration substitution after the EPM testing (see below). For C2, drug concentrations (1 mg/mL and 100 mg/mL) were altered for each session, in a counter-balanced order and rats were exposed to each concentration once. Mean rewards obtained across the last two two-hour self-administration sessions conducted during the EPM testing phase were used as the 50 mg/mL comparison, since these were the last consecutive sessions prior to the substitution experiments that were not conducted after any breaks in testing; weekend or longer breaks imposed by holidays frequently increased responding on the subsequent day in this cohort. Dependent variables for the concentration substitution experiment were the number of reinforcers obtained and the number of active lever presses during the time-outs.

#### 2.4.4 Naloxone challenge

Rats in C2 were returned to self-administration using the training concentration (50 mg/mL in PG vehicle) for one session following substitution experiments. Naloxone challenges (0.0, 0.03, 0.3 and 1.0 mg/kg, i.p.) were then performed over the next four sessions via injection five minutes prior to the session in a counter-balanced order. The sessions lasted 120 minutes to match the ongoing training conditions, but the data were analyzed in 15-minute bins across the session, due to the rapid pharmacokinetics of naloxone. Baseline was calculated as the mean rewards obtained across the last two two-hour self-administration sessions conducted during the EPM testing, segmented into 15-minute bins.

### 2.5 Elevated plus maze test (EPM)

The elevated plus maze apparatus had two opposing open arms and two opposing closed arms perpendicular to the open arms. The arms each measured 50 cm in length and 10 cm in width. The walls of the two closed arms measured 40 cm in height. The apparatus was raised 50 cm from the floor by four legs, one leg located towards the end of each arm. The test was conducted with only a single dim light from a lamp aimed at a wall perpendicular to the open arms. The arms and walls of the apparatus were cleaned thoroughly with an alcohol solution prior to the testing of each animal.

To investigate anxiety-like effects of drug discontinuation, rats in C2 were permitted to self-administer heroin (50 mg/mL) vapor (as described in acquisition methods) for two 2-hour sessions and one 1-hour session on three sequential days; the latter was immediately followed by the first EPM trial. The one-hour session duration was chosen to minimize the time between the initial loading phase, where much of the exposure takes place, and testing. The following day rats underwent a two-hour self-administration session, and the second EPM test was performed 24 hours after completion of this session. Three more two-hour self-administration sessions were completed over three days and the third EPM test was conducted 48 hours after completion of the last session. Rats were motion-tracked during testing, and Open/Closed arm time and entries were analyzed using AnyMaze software (Stoelting, Wood Dale, IL). Duration of EPM testing was set to 300 seconds. The training concentration (50 mg/mL heroin in PG) was used in this study and the starting time of self-administration sessions were offset across individuals to control for time gaps between the end of the session and the beginning of EPM testing.

### 2.6 Data analysis

Data were analyzed by Analysis of Variance (ANOVA) with repeated measures factors for Session in acquisition, Time after exposure in EPM experiments and in nociception assays, and for Concentration or Dose in substitution and naloxone experiments; a paired t-test was used when only two levels of a single factor were included in an analysis. Pre- and post-session tail-withdrawal data collected from rats in C2 following a 30-minute heroin vapor self-administration session were analyzed by paired t-test, and strength of association between tail-withdrawal latency and reinforcers obtained was determined by Pearson correlation. A median split analysis of the acquisition data was planned in advance based on pilot work with the inhalation model, as well as significant differences observed in intravenous self-administration studies (Creehan et al., 2015; Nguyen et al., 2018; Vandewater et al., 2015). Median splits were determined within each Cohort by ranking individuals on mean number of reinforcers obtained across the 10 sessions of acquisition. The upper half is termed High Responder and the lower half is termed Low Responder. Significant effects were followed with post-hoc analysis using the Holm-Sidak procedure. A criterion of p<0.05 was used to infer significant effects. Analysis were performed using Prism 8 for Windows (GraphPad Software, Inc, San Diego, CA).

## 3. Results

### 3.1 Nociception assay

Non-contingent administration of heroin or methadone via injection or vapor delivery increased tail-withdrawal latency (**Figure 1**). Preliminary analysis confirmed there was no significant sex-difference, or interaction of drug conditions with sex, thus the data were collapsed across sex for final presentation and subsequent analysis. The ANOVA for the injection study confirmed significant effects of Drug Condition (F (4, 60) = 45.06; collapsed across sex for final presentation and subsequent analysis. The ANOVA for the injection study confirmed significant effects of Drug Condition (F (4, 60) = 45.06;p<0.0001), of Time post-injection (F (3, 45) = 152.4; p<0.0001) and of the interaction of Time and Drug Condition (F (12, 180) = 32.73; p<0.0001) on tail-withdrawal latency. Post-hoc analysis confirmed that within the 35-minute timepoint, latencies produced by s.c. injection of oxycodone 2 mg/kg (p<0.005), oxycodone 4 mg/kg (p<0.0001), methadone 1 mg/kg (p<0.0001), and heroin 1 mg/kg (p<0.0001) were significantly longer compared with the vehicle group. Post-hoc analysis additionally confirmed that at the 35-minute timepoint, the oxycodone 4 mg/kg group had significantly longer latencies compared with the heroin 1 mg/kg group (p<0.05), and that latencies produced by oxycodone 4 mg/kg (p<0.0001), methadone 4 mg/kg (p<0.0001), and heroin 1 mg/kg(p<0.0001) were significantly longer compared with those from the oxycodone 2 mg/kg group. At the 60-minute timepoint, oxycodone 4 mg/kg (p<0.0001), methadone 4 mg/kg (p<0.0001), and heroin 1 mg/kg (p<0.005) produced significantly longer latencies compared with vehicle. Compared with oxycodone 2 mg/kg, oxycodone 4 mg/kg(p<0.0001), methadone 4 mg/kg (p<0.0001), and heroin 1 mg/kg (p<0.05)produced significantly longer latencies. Also within the 60-minute timepoint, oxycodone 4 mg/kg produced longer latencies compared with methadone 4 mg/kg (p<0.0001) and with heroin 1 mg/kg (p<0.0001), and methadone 4 mg/kg produced longer latencies compared with heroin 1 mg/kg (p<0.005). At 90 minutes, methadone 4 mg/kg produced longer latencies compared with vehicle (p<0.01) and with heroin 1 mg/kg (p<0.05). No differences were confirmed by post-hoc analysis at the 120-minute measurement. Similarly, the ANOVA for the inhalation study confirmed significant effects of Drug Condition (F (3, 45) = 8.194; p<0.001), of Time after the start of inhalation (F (3, 45) = 32.19; p<0.0001) and of the interaction of Time and Drug Condition (F (9, 135) = 10.72; p<0.0001) on tail-withdrawal latency. Within the 35-minute timepoint, the post-hoc analysis confirmed that latencies from the heroin 100 mg/mL (p<0.0001) and methadone 100 mg/mL (p<0.0001) groups were significantly elevated compared with those from the PG group. Additionally, within the 35-minute timepoint, a significant increase in withdrawal latencies was determined in the heroin 100 mg/mL (p<0.0001) and methadone 100 mg/mL (p<0.0001) groups compared with the oxycodone 100 mg/mL group. Within the 60-minute timepoint, Heroin 100 mg/mL (p<0.05) and methadone 100 mg/mL (p<0.0001) were significantly elevated compared with the PG group. Within the 60-minute timepoint, methadone 100 mg/mL produced significantly longer latencies compared with oxycodone 100 mg/mL (p<0.0001). No differences were confirmed within the 90- or 120-minute timepoints.

**Figure 1:**
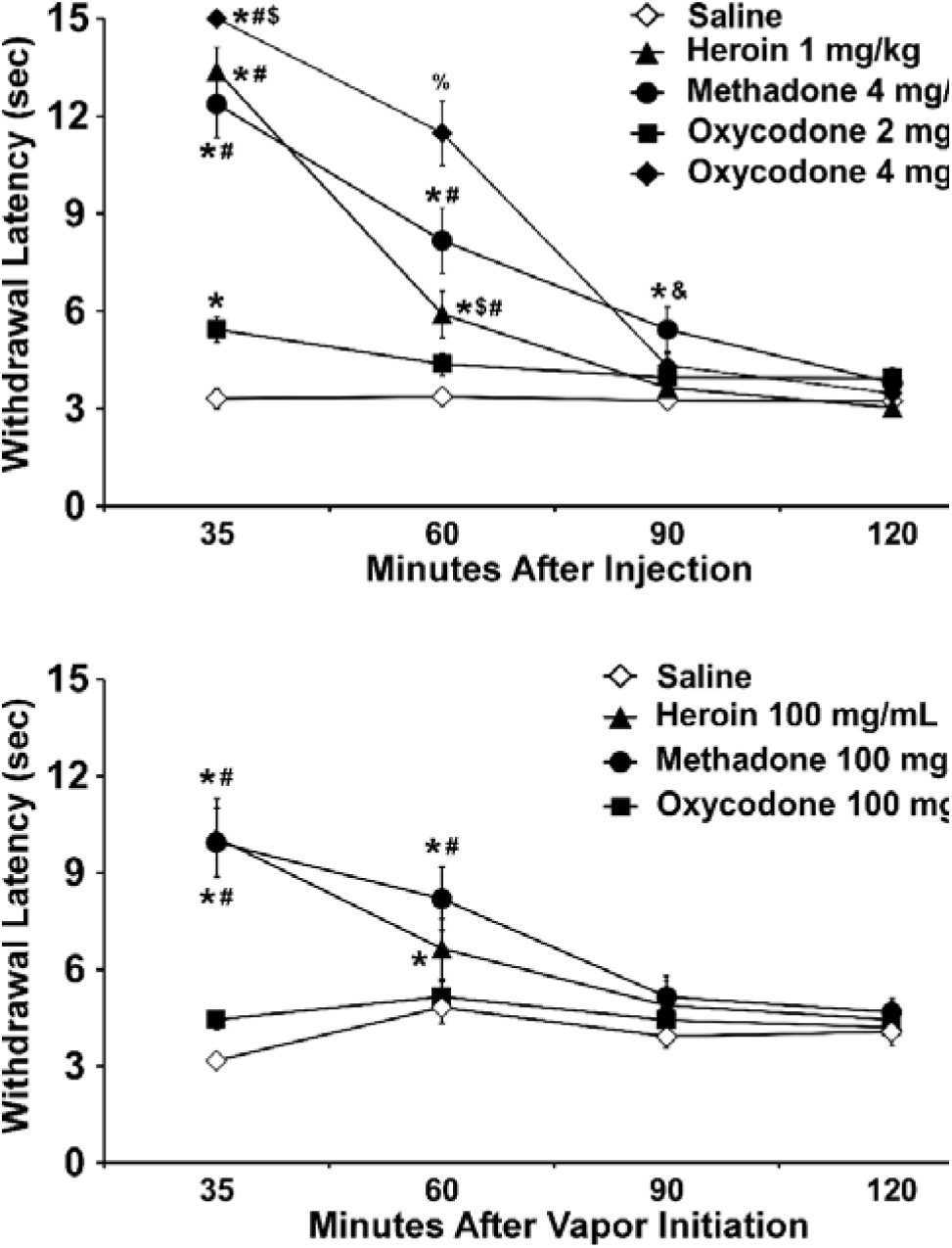
Mean (N=16; ±SEM) tail-withdrawal latencies following (top) s.c. injection or (bottom) inhaled routes of administration for heroin, oxycodone or methadone. A significant difference from vehicle (Saline or PG, respectively) at a given time point is indicated with *, from oxycodone (2 mg/kg or 100 mg/mL) with #, from methadone with $, from Heroin with &, and from all other treatment conditions with %.

### 3.2 Acquisition of self-administration of vaporized heroin

Rats were trained to voluntarily lever-press for contingent vapor deliveries, and consequently exposed themselves to approximately 5-10 minutes of heroin vapor inhalation during the initial 10 sessions (**Figure 2A**). Analysis of reinforcers, by median split, for the FR(1) sessions confirmed a main effect of Group (F(1,14)=8.972, p<0.01; **Figure 2C**), with High Responders (HR) self-administering more reinforcers compared with Low Responders (LR); however, no effect of Session was confirmed (**Figure 2A**). Analysis of reinforcers obtained during the subsequent three FR(2) sessions by median split (**Figure 2C**) did not confirm a significant effect of Group, of Session or of the interaction (p=0.056) despite a recovery in the more-preferring HR rats in Session 13. Analysis of active lever discrimination for the FR(1) sessions confirmed only a main effect of Session(F(9,126)=4.301, p<0.0001; **Figure 2B,D**). Post-hoc analysis confirmed that animals exhibited higher active lever discrimination on sessions 8 (p<0.05) and 10 (p<0.05) compared with session 1, and higher discrimination on session 4 (p<0.05), 8 (p<0.005), and 10 (p<0.005) compared with session 2. No significant effects were confirmed for active lever discrimination for the FR(2) sessions.

**Figure 2:**
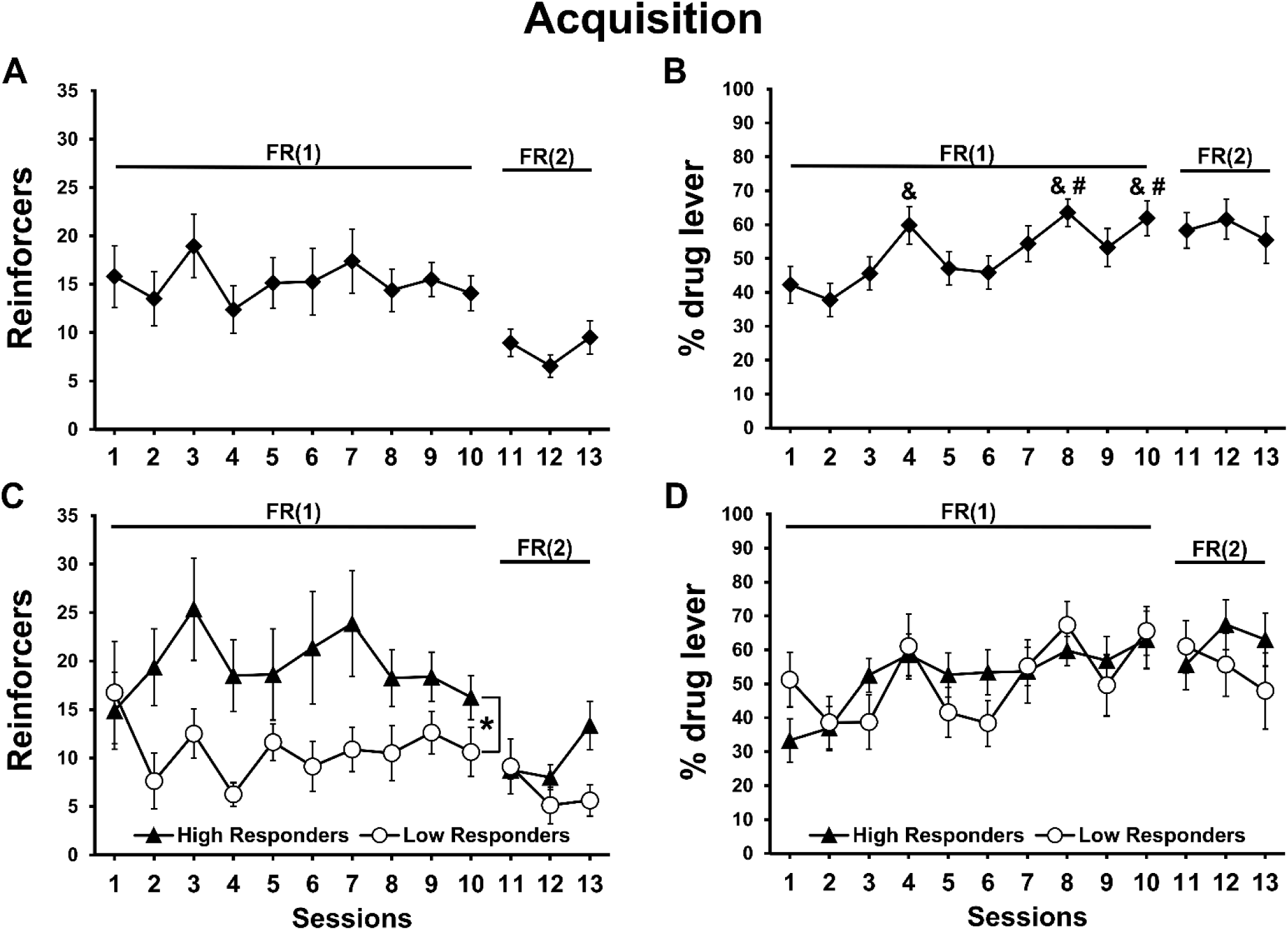
Mean (N=16; ±SEM) A) reinforcers obtained (30 sec epochs of vapor delivery), and B) lever discrimination ratios exhibited, for one-hour heroin vapor self-administration sessions in female rats. Data are presented for High and Low responder sub-groups in panels C and D. Significant differences between HR and LR groups are indicated by *. Significant differences from session 1 and session 2 are indicated by # and &, respectively.

### 3.3 Concentration substitution

Changes in the heroin concentration in the vapor solutions altered the drug seeking behavior of the rats (**Figure 3**). HR (N=8) and LR (N=8) sub-groups were analyzed separately for these experiments. Within the HR group, a significant main effect of Concentration was confirmed for reinforcers obtained (F(2,14)=6.462, p<0.05; **Figure 3A**). Post-hoc analysis confirmed a significant increase in reinforcers obtained for the 1 mg/mL heroin concentration compared with 100 mg/mL heroin concentrations (p<0.01). No main effect of Concentration was confirmed within the HR group for time-out drug-associated lever responding (p = 0.05; **Figure 3C**). The mean number of reinforcers obtained by the LR rats did not differ across concentrations (**Figure 3B**) and time-out responses were not statistically different (p = 0.076; **Figure 3D**).

**Figure 3:**
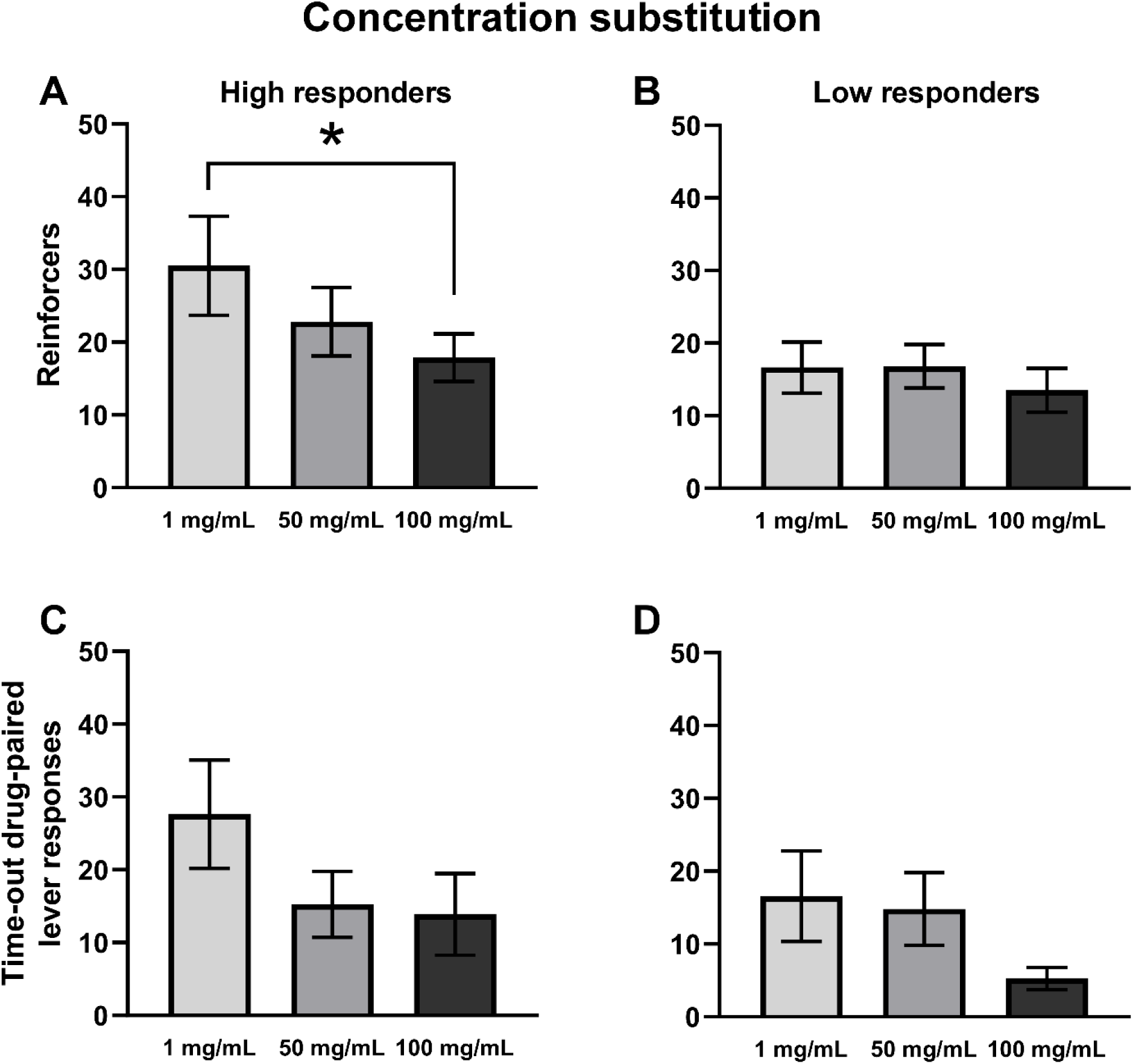
Mean (±SEM) reinforcer deliveries (A, B) and time-out drug-associated lever responding (C, D) for High (A, C) and Low (B, D) Responder sub-groups. A significant difference between concentrations is indicated with *.

### 3.4 Naloxone challenge

Pre-treatment with the mu opioid receptor antagonist increased the number of reinforcers obtained by the HR rats, in a dose-dependent manner (**Figure 4**).

**Figure 4:**
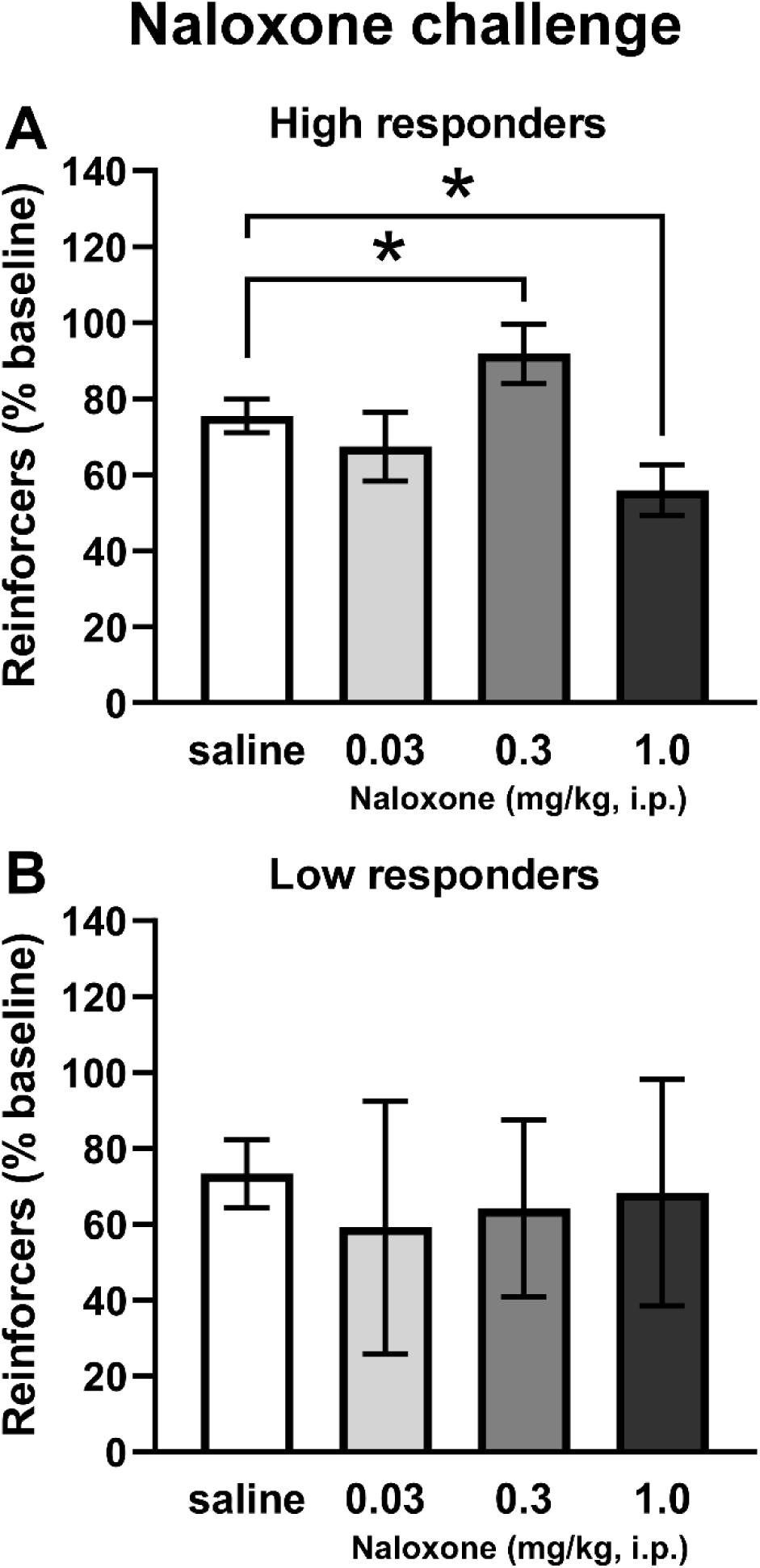
Mean (±SEM) reinforcers obtained by HR (A; N=4) and LR (B; N=4) sub-groups of Cohort 2 following i.p. naloxone injection. A significant difference between pre-treatment conditions is indicated with *.

Reinforcers obtained in the first 15 minutes of the session by HR and LR rats (N=4 per treatment condition) were analyzed separately for these experiments and the data are presented as % of baseline to illustrate an apparent effect of the pre-session injection protocol. The ANOVA confirmed a significant effect of Dose (F(3,9)=12.81, p<0.01) on reinforcers obtained by the HR rats (**Figure 4A**). The post-hoc analysis confirmed a significant increase in the 0.3 mg/kg dose condition (p<0.05), and a decrease in the 1.0 mg/kg dose condition (p<0.05), in comparison with saline pre-treatment. No significant effects of pre-treatment condition were confirmed in the LR group (**Figure 4B**).

### 3.5 Anti-nociceptive effects of self-administered heroin

Tail-withdrawal latencies were recorded prior to any drug exposure, and before and after the eighth self-administration session for rats in both cohorts (**Figure 5A**). There was no significant difference in tail-withdrawal latency between HR and LR groups, thus the data were collapsed across Responder groups. The analysis confirmed a significant effect of Timepoint (F(2,30)=15.61, p<0.0001) on tail-withdrawal. Post-hoc analysis confirmed that tail-withdrawal latency was significantly longer post-session compared with either baseline (p<0.0001) or pre-session (p<0.0005) measurements. No differences were confirmed between baseline and pre-session time points. A second tail-withdrawal test was performed on rats in C2 prior to and after the 20^th^ self-administration session, which was shortened to 30 minutes duration to constrain variability in intake pattern. Because no effect of preference sub-group was confirmed in the first tail-withdrawal assay, these data were also analyzed as one group. A paired t test confirmed a significant difference between the pre- and post-session measurements (t(7)=2.921, p<0.05; **Figure 5B**); post-session withdrawal latencies were significantly higher compared with pre-session. A Pearson correlation also confirmed a significant positive association between the number of reinforcers obtained during the self-administration session and tail-withdrawal latency in the 30-minute experiment (r=0.8008, p<0.05; **Figure 5C**).Analysis of tail-withdrawal latency data from a separate group of male rats that were permitted to self-administer heroin intravenously for 30 minutes (**Figure 5D**) confirmed a significant main effect of time of assessment (F(1,15)=8.012, p≤0.01); post-session withdrawal latencies were significantly higher after heroin IVSA compared with pre-session latencies.

**Figure 5:**
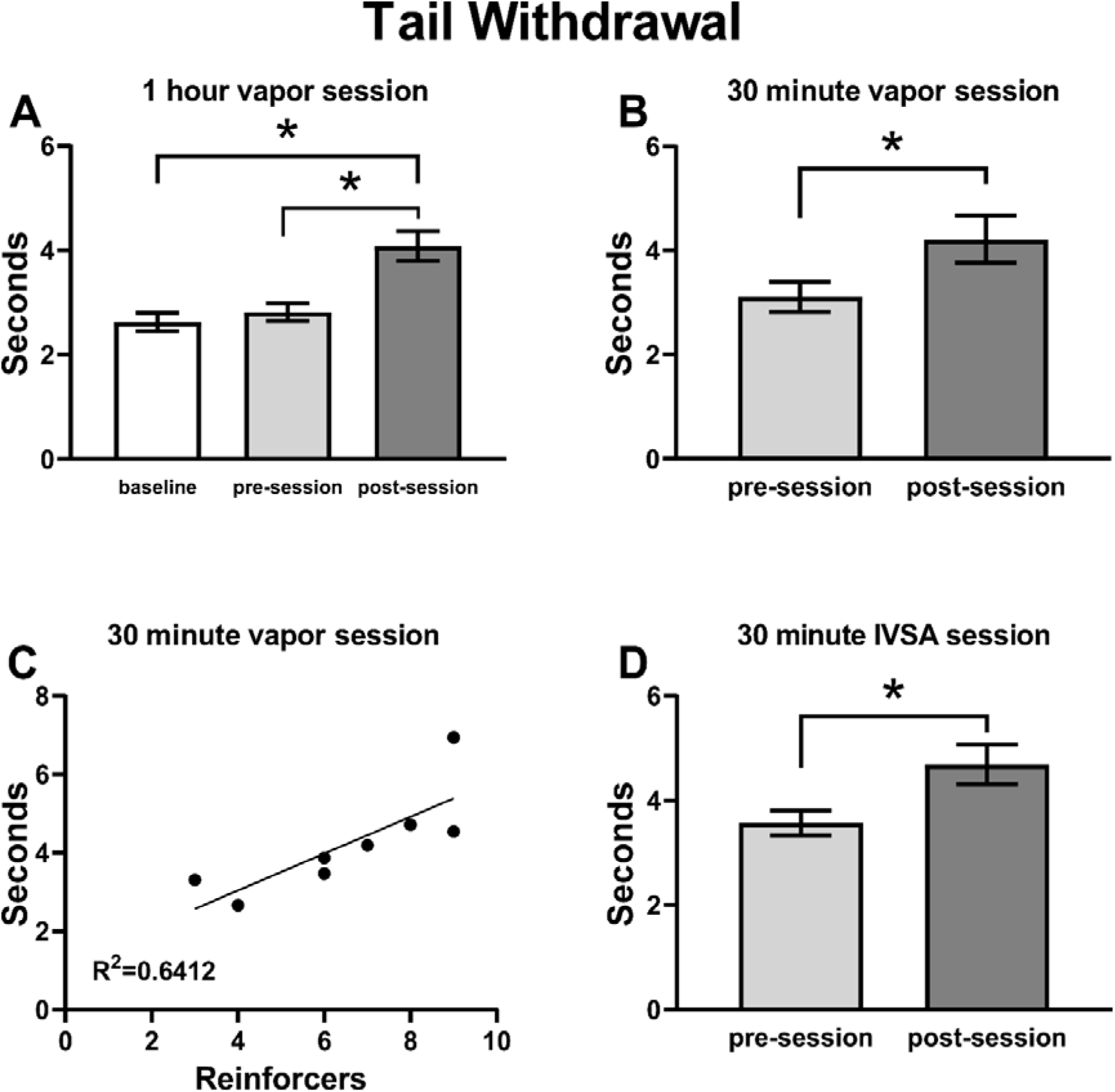
Mean (±SEM) tail withdrawal latency for A) Cohorts 1 and 2 (N=16); B) Cohort 2 (N=8) rats before and after a 30 min heroin vapor session; C) Correlation of individual (N=8) withdrawal latency with the number of reinforcers obtained in the prior self-administration session for the 30 min vapor inhalation experiment; D) Male Wistar (N=17) rats trained in intravenous heroin self-administration and assessed before and after the session. A significant difference between assessments is indicated with *.

### 3.6 Elevated plus maze

The time rats spent in the open and closed arms of the EPM varied depending on the interval of time since last drug access (**Figure 6**). The statistical analysis of theEPM behavior confirmed a significant effect of the post-session interval for time spent in the closed arms (F(2,14)=6.233, p<0.05; **Figure 6A**) and time spent in the open arms (F(2,14)=7.030, p<0.01; **Figure 6B**) of the maze. The post-hoc analysis confirmed an increase in time spent in the closed arms at the 48-hour time point compared with the immediately post-session assessment (p<0.05) as well as a significant decrease in the time spent in the open arms at both the 24-(p<0.05) and 48-hour (p<0.01) post-session time points compared with the immediate post-session time point. Analysis of arm entries also confirmed significant effects of drug discontinuation time for entries into the open arms (F(2, 14)=7.366, p<0.01; **Figure 6D**) and the post-hoc test confirmed fewer mean entries 24-(p<0.05) and 48-hour (p<0.05) after last drug availability compared with the immediate post-session assessment. No effects of time since last drug availability were confirmed for closed arm entries (**Figure 6C**).

**Figure 6:**
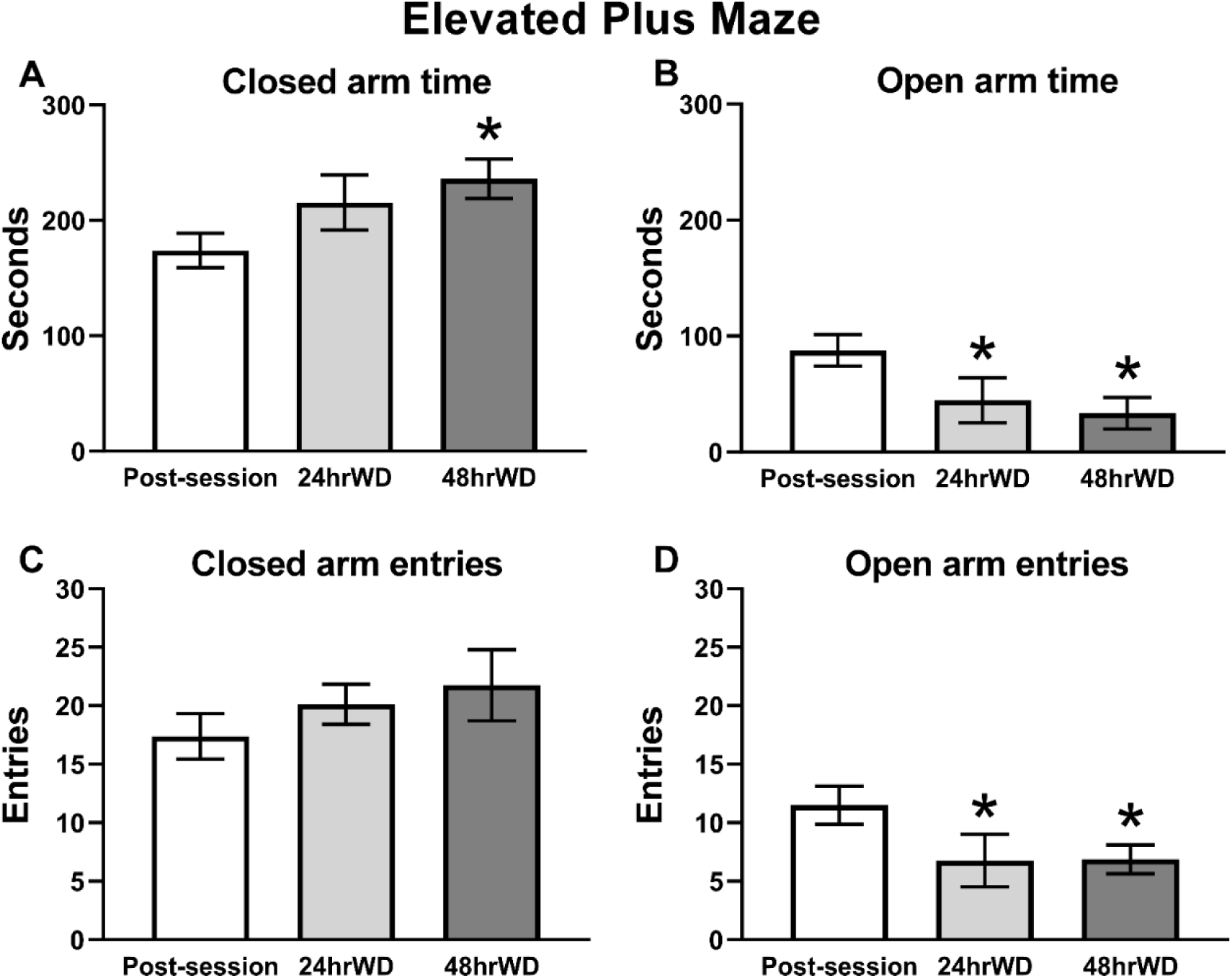
Mean (N=8; ±SEM) time spent in A) closed and B) open arms, and number of entries into the C) closed and D) open arms of the elevated plus maze. A significant difference from the post-session assessment, is indicated with *.

## 4. Discussion

In the present study, heroin vapor generated by an Electronic Drug Delivery System (EDDS; “e-cigarette”) reinforced operant responding in a manner that is consistent with heroin self-administration. This extends prior results using extended daily access sessions (12 h) and the high potency opioid sufentanil (Vendruscolo et al., 2018) to the less-potent but more commonly abused opioid heroin, to short-access sessions and to female rats. The first major finding of this study was the demonstration that non-contingent delivery of heroin and methadone by EDDS technology produced significant analgesia in rats. Thus, this approach can be useful for the assessment of a wide range of effects of several common opioids. The second major finding of this study was that the higher preference rats met two key criteria for self-administration, i.e. they compensated for changes in the heroin concentration in the vapor with changes in behavioral responding, and they similarly compensated for the effects of a moderate dose of the opioid receptor antagonist naloxone. Furthermore, acute intoxication via vapor self-administration was shown to produce anti-nociception comparable to that produced by intravenous self-administration of heroin. This outcome confirms the similarity of the self-determined heroin intake level by the two routes of administration, and provides additional evidence supporting interpretation of the vapor inhalation behavior as heroin self-administration. The fact that individual differences in anti-nociception correlated with the number of vapor deliveries obtained in the prior self-administration session provides converging evidence on the function of the model. Lastly, anxiety-like behavior, assessed in an EPM, increased following 24 and 48 hours of drug discontinuation. This pattern suggests that withdrawal-like effects develop shortly after access to heroin vapor is withheld, similar to what is observed after access is withheld in an intravenous self-administration model (Barbier et al., 2013).

In the concentration-substitution experiment the High Responder (HR) rats self-administered a mean of ∼20 vapor deliveries at the training concentration and were sensitive to an increase in the heroin concentration since they decreased the number of obtained reinforcers to compensate. The time-out responding of HR rats on the drug-associated lever also increased when the concentration was lowered, suggestive of enhanced drug-seeking associated with the lower available concentration of heroin. The Low Responder (LR) rats did not consistently increase responding when the concentration was decreased relative to the training dose. But they did, however, decrease their time out responding at the higher concentration of heroin. This pattern is consistent with an interpretation that this subgroup was seeking opioid intoxication, but was perhaps more sensitive to aversive properties of higher levels of intoxication. If so, this could therefore be viewed as evidence of overall purposeful, drug-motivated behavior on the part of the lower-preferring individuals. This may explain why they continued to respond throughout the entire course of study, i.e., no rats in the group ever completely extinguished their responding.

Treatment with an antagonist such as naloxone can dose-dependently alter responding for opioid drugs, enhancing response rates at lower doses and suppressing responding as the dose is increased. For example, naloxone given at 1 mg/kg suppressed responding for aerosolized sufentanil in rats (Jaffe et al., 1989) and smoked heroin in rhesus monkeys (Mattox and Carroll, 1996), and increased responding for intravenously administered heroin in rats when given at doses ranging from 0.003 to 0.3 mg/kg (Carrera et al., 1999; Chen et al., 2006). For the higher preference subgroup in this study, naloxone altered vapor self-administration in an inverted U dose-effect pattern suggesting that the intermediate dose attenuated heroin reward whereas the higher dose may have prevented it under these inhalation conditions. Results were not as consistent for the LR subgroup, however three of the four increased their responding, compared with saline pre-treatment, following at least one naloxone dose. This aspect of the study further supports the conclusion that rats were responding to self-administer an individually determined dose of heroin via inhalation.

Tail-withdrawal latencies were reliably increased in the nociception assays conducted after a self-administration session, which is consistent with the established anti-nociceptive properties of heroin. The effect size was not large compared with the effects of larger parenteral injection of heroin (Tasker and Nakatsu, 1984), methadone (Holtman and Wala, 2007) or oxycodone (Nguyen et al., 2019) as reported in prior studies and in our validation (Figure 1, top), or a non-contingent delivery of methadone or heroin vapor (generated from 100 mg/mL) for 30 minutes (Figure 1, bottom). Some degree of tolerance due to the repeated self-administration might be expected, for example, tolerance to anti-nociceptive effects of a 1 mg/kg, s.c. dose of oxycodone is induced by an interval of intravenous oxycodone self-administration, as we previously showed (Nguyen et al., 2018). Nevertheless, this outcome likely points out that self-administered doses of psychoactive drugs are often far lower than those which produce maximal effects in non-contingent assays of other responses, *in vivo*. Most importantly, we show here that the magnitude of anti-nociception produced by self-selected dosing of inhaled heroin is identical to that produced by a self-selected dose of i.v. heroin. This supports the conclusion that rats are self-administering to a roughly similar degree of heroin intoxication in an inhalation session as they do when *intravenous* infusions are made contingent upon a lever press.

The elevated plus-maze test and similar approaches, such as the elevated zero maze test (Abdulla et al., 2020; Braun et al., 2011; Shepherd et al., 1994; Sprowles et al., 2016), are well established behavioral assays for assessing anxiety-like behavior (Handley and Mithani, 1984; Montgomery, 1955; Pellow et al., 1985). In the present study, rats exhibited signs of increasing anxiety-like behavior across 24 h and 48 h of drug abstinence in this short-access (2 h) model, using the elevated plus-maze approach. This is consistent with results for discontinuation from intravenous self-administration of heroin under 1 h (Barbier et al., 2013; Schlosburg et al., 2013) or 4 h access conditions (Lou et al., 2014). As a minor caveat, in this study, EPM testing was performed over repeated trials for each individual. A concern of this approach is the impact that repeated exposure may have on key behavioral measures, such as open arm time. Although stability has been reported in non-treated rats (Schrader et al., 2018; Tucci et al., 2002), in other cases it has been reported that closed-arm time and/or entries may increase or decrease with repeated testing under un-drugged conditions (Bertoglio and Carobrez, 2002; Roy et al., 2009). Furthermore, effects of previous maze experience under drugged conditions on subsequent sessions (Tucci et al., 2002) and of non-drugged maze experience on drug efficacy during subsequent sessions (File, 1990) have been reported. The interplay between experience, behavioral drug effects, and affective states was not the focus of this study, however, and the extent to which the design affected withdrawal expression cannot be precisely determined from the present study. Importantly, however, the findings here are in agreement with previous studies assessing withdrawal-related anxiety-like behavior in opioid-treated animals using the EPM.

One apparent difference in the vapor self-administration model, relative to conventional IVSA criteria for self-administration, is the relatively low ratio of responses on the drug-associated versus the non-associated lever. Similar results have been reported in mice responding for fentanyl vapor during 1 hour sessions at a FR(1) schedule of reinforcement (Moussawi et al., 2020). Relatedly, the rats in the present study also tended to make responses on the drug-associated lever during the time-out interval. This result in our study mirrors similar patterns reported by Freels and colleagues and by Vendruscolo and colleagues (Freels et al., 2020; Vendruscolo et al., 2018) for the self-administration of cannabis and sufentanil vapor, respectively. Thus, these patterns may be commonalities of vapor self-administration, or consequences of the methodological approaches that have been tried so far. Another limitation is the distribution of individual differences. One recent vapor self-administration study in mice similarly reported a number of animals that failed to reach stable responding for nicotine vapor (Cooper et al., 2020), echoing the Low Responder subset in this investigation. Similar distributional differences in responding have also been reported for the acquisition of intravenous self-administration of 3,4-methylenedioxymethamphetamine (Schenk et al., 2007; Vandewater et al., 2015) and even for cocaine if a subthreshold training dose is used (Smith and Pitts, 2011), thus this is not entirely unexpected. Further development of these inhalation models will undoubtedly lend greater insight into approaches that diminish or enhance intragroup variability and whether it is preferable to use acquisition screening criteria or sub-population analyses in various experimental designs. Additionally, it should be noted that although oxycodone vapor inhalation did not work in the present anti-nociception experiment using the Pro-tank 3 EDDS device, we have shown efficacy of oxycodone alone, and in synergy with Δ^9^-tetrahydrocannabinol, using the SMOK Baby Beast canisters (Nguyen et al., 2019). Benefits of EDDS delivery of this popular prescription opioid may therefore also be investigated, but this is a reminder that EDDS device characteristics may be critical to the success of some experimental approaches.

There are undoubtedly many additional variants of the inhalation self-administration approach to explore which may turn out to have advantages and disadvantages relative to the current model. This work should be viewed as an initial demonstration of feasibility and a departure point for examining other variations in approach, which may have improved utility across a range of experimental goals. Training doses, vapor dwell time, chamber size and other technical variants may influence self-administration behavior and should be investigated in follow-up studies. Additional questions of future interest would involve understanding how sex, rat strain or developmental age may contribute to heroin vapor self-administration.

## 5 Conclusions

These studies show that self-administration of heroin by vapor inhalation in rats using Electronic Drug Delivery System (“e-cigarette”) technology can be produced in short-access models. This extends prior observation of the establishment of vapor inhalation of sufentanil (Vendruscolo et al., 2018) using extended (12 hour) daily access sessions, fentanyl in short or long access sessions (Moussawi et al., 2020), cannabis extracts (Freels et al., 2020) or nicotine (Cooper et al., 2020).

## Acknowledgements

The authors are grateful to Sophia A. Vandewater for assistance with blinding of the initial anti-nociception experiment, and to Maury Cole of La Jolla Alcohol Research, Inc, La Jolla, CA, USA for design of the vapor self-administration equipment. These data were initially made available in pre-print form (Gutierrez et al., 2020b).

## Funding

This work was supported by USPHS grants T32 AA007456 (Gutierrez, Fellow), R01 DA035281 (Taffe, PI), K99 DA047413 (Nguyen, PI), R44 DA041967 (Cole, PI), by a UCSD Chancellor’s Post-doctoral Fellowship Program (Gutierrez, Fellow), and by the Tobacco-Related Disease Research program of California (T31IP1832). The NIH/NIDA and the TRDRP had no direct influence on the design, conduct, analysis or decision to publish findings. La Jolla Alcohol Research Inc. (to which the R44 was awarded) likewise did not influence the study designs, the data analysis or the decision to publish findings.

## Author Contributions

AG, JDN and MAT designed the studies. AG, KMC and JDN performed the research and conducted initial data analysis. AG conducted statistical analysis of data, created figures and AG and MAT wrote the initial draft of the paper. All authors approved of the submitted version of the manuscript.

## Notes

### Competing Interest Statement

The authors have declared no competing interest.

### Summary of Updates

This version has provided additional specification of a few experimental details, expansion of the description of the post-hoc exploration (including p values), revision of the caveats on the elevated plus maze experimental design and a broadening of the discussion of additional findings with vapor exposure models.

